# The mandibular gland in *Nasonia vitripennis* (Hymenoptera: Pteromalidae)

**DOI:** 10.1101/006569

**Authors:** István Mikó, Andrew R. Deans

## Abstract

The mandibular gland of *Nasonia vitripennis* (Hymenoptera: Pteromalidae) is visualized for the first time, using Confocal Laser Scanning Microscopy and dissection. The gland was previously hypothesized to exist, based on observations of the wasp’s courtship behaviors, but its presence had never been confirmed.

## INTRODUCTION

*Nasonia vitripennis* (Hymenoptera: Pteromalidae; Jewel Wasp) is the most researched parasitoid in the model organism/genomic world. The reasons, in part, are their easy handling, the availability of cultures, the small body size, and the species’ very short developmental period. While it attracts genomic people to study gene function and to test findings from *Drosophila*, these minute wasps are really interesting in other ways as well. They are among the few organisms where males attract virgin females. They do this with a pheromone—(4R,5R)- and (4R,5S)-5-hydroxy-4-decanolide (HDL)—that is a product of the rectal gland (Steiner and Ruther, 2009).

There is another, perhaps even more exciting but certainly less understood behavioral aspect of *Nasonia* males, related to the activity of a second pheromone producing gland in males. Van den Assem and colleagues have performed several experiments on these small wasps (e.g., Assem et al. (1980)) that, as far as we know, have never ever been repeated and thus their findings never confirmed. (Barrass, 1960, p. 191) first described a unique courtship position for male *N. vitripennis*:

> The male’s head is above and between the female’s erect antennae and its mandibles are very near to the proximal region of the female’s flagella. In this position (Figures I and 2A) the antennae are held above and anterior to those of the female. Following this the male’s head moves downwards slightly (Figure 2B), and then it is raised so that the mouthparts are near to the distal region of the female’s flagella.

The courtship head movement, later named as head nodding by van den Assem and his colleagues is a unique behavior of *Nasonia* males that might actually improve the discharge of an enigmatic sexual pheromone, an aphrodisiac. Although uncommon, aphrodisiacs do occur in numerous insect taxa. They have been reported from many Lepidoptera, Dictyoptera, Orthoptera, and Heteroptera (Butler, 1967) and are usually produced by males for preparing the female for copulation, after the couple has been brought together by olfactory sex attractants and/or other means.

van den Assem and colleagues hypothesized that a pheromone is produced by the male during courtship that has aphrodisiac effect, and that perhaps it is related to the nodding movement of the head. In the their experiments, the researchers combined the following treatments: (1) sealed the mouthparts with superglue, (2) removed the abdomen and left the propodeal foramen open, the propodeal foramen is the opening that serves as the passage for nerves, trachea, alimentary tract, some muscles and hemolymph between the meso- and metasoma, (3) removed the abdomen and sealed the propodeal foramen with superglue.

Maimed males without sealed propodeal foramena were unable to nod their heads or extrude their mouthparts but they gained back the ability to perform these behavior if the sealing of their propodeal foramen took place shortly after the “amputation” of the metasoma. Normal males with sealed mouthparts (nodding is possible but mouthpart extrusion and mandible movement not) and maimed males with unsealed propodeal foramen were not able to provoke receptivity in females, whereas control (untouched) males and males with sealed propodeal foramen were able to increase the “sexual desire” of female wasps.

The results of these quite superglue intensive experiences clearly shows that the male *Nasonia vitripennis* specimen producing a pheromone somewhere around the mouthparts and that the head nodding and mouthparts extrusion are important factors for the aphrodisiac effect. The authors hypothesized that: (1) There has to be an intricate system operating with pressure change of the hemolymph required for head nodding and mouthparts protrusion and thus gland release. And (2) the mandibular gland is the source of the enigmatic aphrodisiac. And that’s it. Two papers published in 1980 and 1981 and not a single follow up! How can it be? The observations of van den Assem *et al*. certainly raised plenty of questions to be answered, but somehow this subject on *Nasonia* aphrodisiac has been sunk into oblivion. Were the hypotheses of van den Assem *et al*., right? What are the compounds of this pheromone? Is it really produced by the mandibular gland? Where is it located then and how is it released? And does *Nasonia vitripennis* even have a mandibular gland?

## METHODS AND MATERIALS

We acquired live *Nasonia* individuals from our colleague (Jack Werren, University of Rochester). CLSM images were taken on glycerin-stored specimens with Zeiss LSM 710 Confocal Microscope. We used an excitation wavelength of 488 and an emission wavelength of 510–680 nm, detected using two channels and visualized separately with two pseudocolors (510–580 nm = green; 580–680 nm = red). To visualizing resilin we used an excitation wavelength of 405 nm and an emission wavelength of 510–680 nm, detected on one channel and visualized with a blue pseudocolor. Specimens were subsequently dissected and studied with an Olympus SZX16 stereomicroscope with SDFPLAPO 2XPFC objective.

## RESULTS AND DISCUSSION

The mandibular gland is visible in Figure 1.

**Figure 1.**
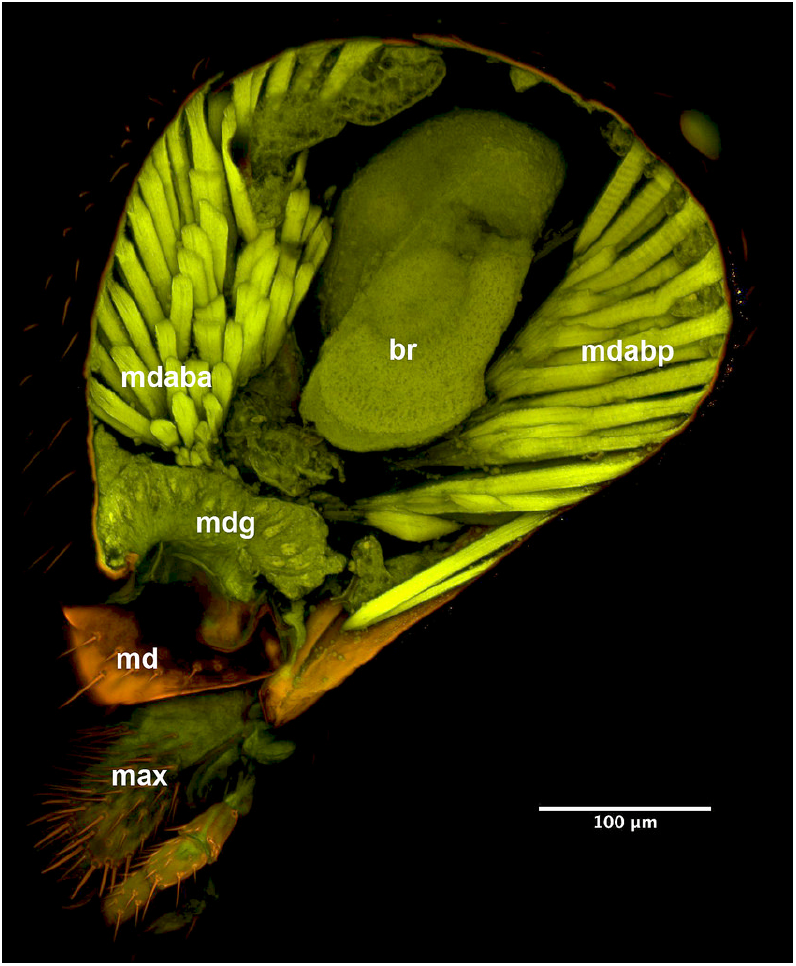
CLSM volume rendered image of *Nasonia vitripennis* male showing the mandibular gland. br = brain, mdaba = anterior branch of mandibular adductor, mdap = posterior branch of mandibular adductor, mgd = mandibular gland, md = mandible, max = maxilla. Photo by István Mikó (CC BY 2.0).

## ACKNOWLEDGMENTS

We thank Jack Werren for supplying us with live individuals. This research was funded by the U. S. National Science Foundation (grants DBI-0850223, DEB-0842289) and by the Phenotype Research Coordination Network (NSF DEB-0956049). The funders had no role in study design, data collection and analysis, decision to publish, or preparation of the manuscript.

